# A single microRNA miR-195 rescues the arrested B cell development induced by EBF1 deficiency

**DOI:** 10.1101/2024.07.26.605276

**Authors:** Y. Miyatake, T. Kamakura, T. Ikawa, R. Yanagiya, R. Kotaki, K. Kameda, R. Koyama-Nasu, K. Okuyama, K. Hirano, H. Hosokawa, K. Hozumi, M. Ohtsuka, T. Kishikawa, C. Shibata, M. Otsuka, R. Maruyama, K. Ando, T. Kurosaki, H. Kawamoto, A. Kotani

## Abstract

Accumulated studies have reported that hematopoietic differentiation was primarily regulated by transcription factors. Early B cell factor 1 (EBF1) is an essential transcription factor for B lymphopoiesis. Contrary to the canonical notion, we found that a single miRNA, miRNA-195 (miR-195) transduction let EBF1 deficient hematopoietic progenitor cells (HPCs) express CD19, carry out V(D)J recombination and class switch recombination, which implied that B cell matured without EBF1. A part of the mechanism was caused by FOXO1 accumulation via inhibition of FOXO1 phosphorylation pathways in which targets of miR-195 are enriched. These results suggested that some miRNA transductions could function as alternatives to transcription factors.

## Introduction

Developmental hierarchy in hematopoiesis has been widely researched, and it is well-known that proper stimulation leads hematopoietic stem cells (HPCs) into B cell lineage. Lineage specification is primarily regulated at the transcriptional level, thus lineage-specific transcription factors are considered to indispensable for differentiation (*1*, *2*). B cell development requires multiple transcription factors, especially Early B cell Factor 1 (EBF1), Paired Box 5 (Pax5), and E2A. Pax5 and E2A are critical transcription factors for early B cell development, but they cannot rescue EBF1-deficient HPCs from failure of B cell lineage commitment (*3*). Conversely, ectopic expression of EBF1 is able to rescue Pax5, E2A, and PU.1 deleted progenitor cells from B lymphopoiesis arrest, and thus EBF1 is considered more potent than the other transcription factors (*2*, *4*, *5*). As the most potent transcription factor, EBF1 is essential for pre-pro-B cell to become pro-B cell; namely, *Ebf1*^-/-^ cell express B220 but is disable to express CD19 (*6*).

MicroRNAs (miRNAs) are small non-coding RNA containing approximately 22 nucleotides that regulate several target protein expressions mediating deadenylation and translation by post-transcriptionally repressing or decaying target messenger RNAs (mRNAs) (*7–10*). Although similar to transcription factor, miRNAs regulate large numbers of target mRNAs and deeply contribute to various cell events, the regulation is mainly required for negative regulation of leaky gene expression and often called as fine-tuning (*11*, *12*). In hematopoiesis, miRNAs are expressed in lineage specific manner and their profiles greatly influence on cell differentiation (*13–15*). Focusing on B cell development, it is revealed that Dicer, a key enzyme of miRNA generation, is essential for pre-to pro-B cell transition (*16*). Individual miRNA is also studied and miR-150 and miR-126 are identified as relational factor to B cell lineage development. miR-150 regulate B cell differentiation by controlling c-Myb expression and miR-126 partially rescues EBF1 deficient B cell lineage commitment by modulating IRS-1 expression (*17*, *18*). Both miRNAs dramatically contributed to B cell development processes, but they were not able to recover B cell development from EBF1 deficiency. Conceived from these vigorous functions of miRNAs on B cell development, in this time, we analyzed ability of miR-195, recently revealed as an important factor for several cell differentiation, on B cell lineage commitment in EBF1 deficient HPCs (*19*, *20*).

## Results

### miR-195 induces B cell character in EBF1 deficient HPCs

To assess the contribution of miR-195 on B cell development, miR-195 was transduced into mouse fetal liver (FL)-derived Lin^-^ c-kit^+^ HPCs and the cells were differentiated to B220 and CD19 expressing pro-B cells with IL7, Flt ligand and SCF on OP9 stroma cells. After 7 days of culture, certain numbers of the cells gradually expressed CD19, and the positive cells was increased by miR-195 transduction (Fig. 1A). This result suggests that miR-195 has ability to shift the HPCs differentiation toward B cells. Next, we attempted to differentiate *Ebf1*^-/-^ FL HPC to B cell with miR-195 transduction (Fig. 1B). As by a previous study (*6*), control *Ebf1*^-/-^ FL HPCs expressed B220 but did not express CD19.

**Fig. 1.**
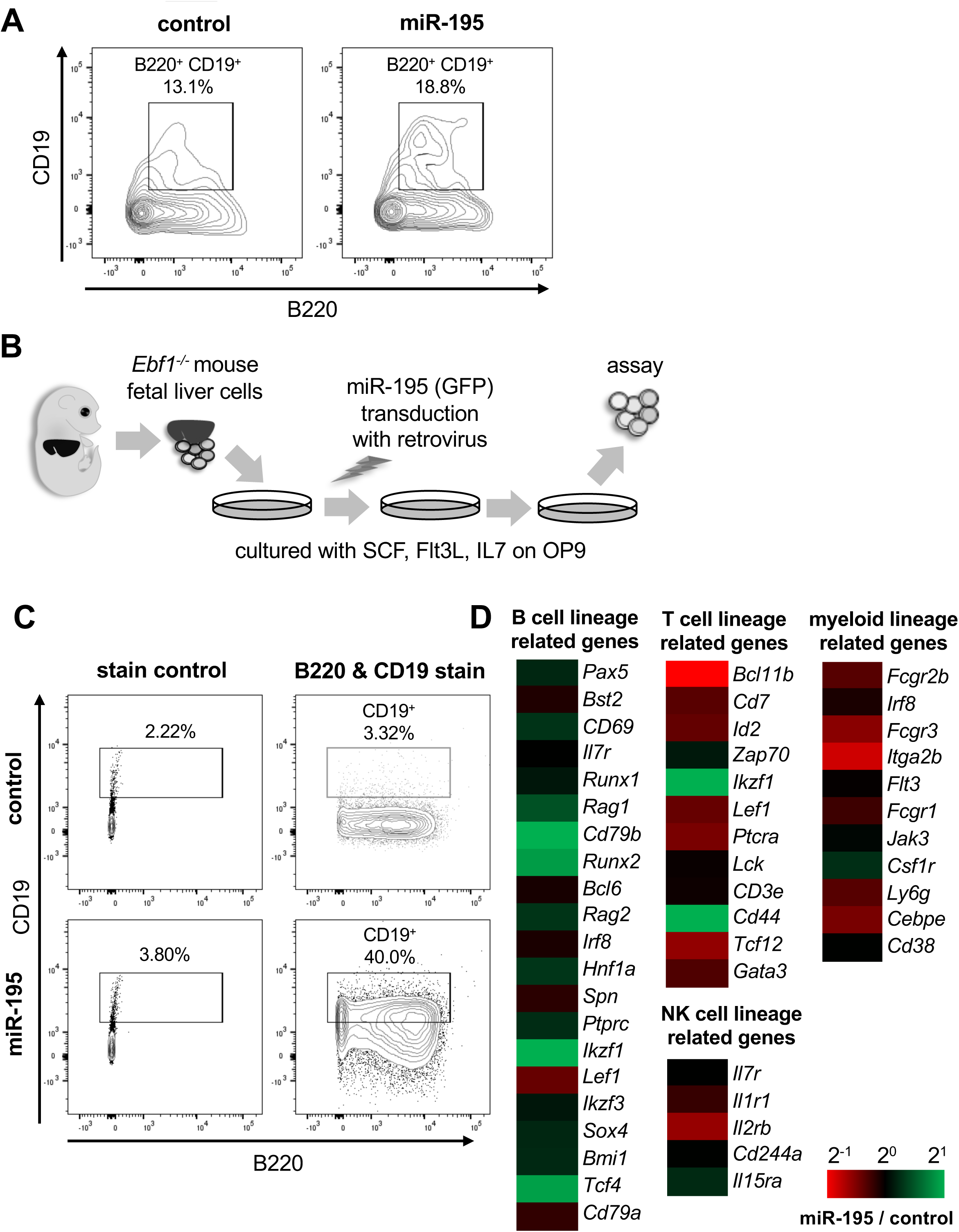
miR-195 promotes HPCs to differentiate into the pro-B cell stage without EBF1. (A) Flow cytometry analysis of control and miR-195-expressing Lin^-^ cells. HPCs from fetal livers of wild-type mice were cultured for 7 days on OP9 with SCF, Flt3-ligand, and IL-7, after infection with control or miR-195 retrovirus. Representative result of control (upper panel) and miR-195 (lower panel) viral infections is shown (*n*=3). (B) Outline of the *in vitro* culture system of *Ebf1***^-/-^** HPCs. (C) Flow cytometry analysis of control and miR-195-expressing *Ebf1***^-/-^**HPCs. Shown data is representative of *n*=3. (D) Microarray analysis of miR-195-expressing *Ebf1***^-/-^** HPCs. Log_2_ fold-changes in the expression levels of genes related to B (left panel), T (middle-upper panel), NK (middle-lower panel), and myeloid (right panel) cell lineages were classified and are shown as colored columns. The analysis was carried out in duplicates.

However, miR-195 transduced *Ebf1*^-/-^ FL HPCs highly expressed CD19 (Fig. 1C). In normal B cell development, CD19 expression follows B220 expression and hence CD19 positive cells show B220 expression as well. Thus, miR-195-transduced *Ebf1*^-/-^ FL HPCs which including B220 negative CD19 positive population may simply reflect up-regulation of CD19 expression, but not B cell development. To exclude this possibility, gene expressions of miR-195-transduced *Ebf1*^-/-^ FL HPCs by cDNA microarray assay were investigated then indicated that miR-195-transduced cells more expressed B cell lineage related genes, e.g., *Pax5, Aicda*, *Rag1*, *Rag2*, *CD79b* and *Runx2*, whereas less expressed T cell and NK cell lineage related genes, including *Gata3, Id2*, *Lck*, *CD3e* and *Il2rb* and also myeloid lineage related genes for example, *Cebpe*, *Ly6g*, *Fcgr1*, *Fcgr2b* and *fcgr3* (Fig. 1D). (1–6) Among the B-lineage transcription factors, *Pax5* and *Erg* were modestly but significantly upregulated (log₂FC ∼1.2 and ∼0.9, respectively) in miR-195-transduced *Ebf1*^−/−^ cells compared to controls. While these changes were moderate, they were consistent across replicates and suggest partial restoration of the B cell transcriptional program. These results suggested that not only CD19 expression but also up-regulation of several B cell developmental factors and down-regulation of other lineage related genes were involved in the promotion of B cell lineage commitment by miR-195.

### EBF1 deficient HPCs were able to commit B cell lineage by transduction of miR-195 with bone marrow niche modification

The ectopic miR-195 expression led *Ebf1*^-/-^ HPCs induced differentiation toward B cell. However, a large part of the miR-195-transduced HPCs expressed CD19 but not B220, which implied that they strayed from the canonical B cell differentiation steps (Fig. 1C). In addition to the inner state, the microenvironment known as niche was also critically involved in hematopoiesis (*21*). Especially in early B cell development, bone marrow niches precisely control the maintenance and differentiation of lineage precursors by cytokines and chemokines (*22*). To explore the development of miR-195-transduced *Ebf1*^-/-^ FL HPCs under bone marrow niches, we engrafted miR-195-transduced *Ebf1*^-/-^ early B cells into NOG and B6RG mice, in which absence of B cell makes the engrafted B cell visible (Fig. 2A). After 7 days, the engrafted cells successfully adapted in the bone marrow. While there was no remarkable change in control cell population, notably, instead of B220 negative CD19 positive cells, the double positive cells were markedly increased in miR-195-transduced *Ebf1*^-/-^ FL early B cells, suggesting that the normal stepwise B cell development occurred (Fig. 2B). In B cell development, most prominent steps after CD19 expression are VDJ recombination and subsequential IgM expression on cell surface. In addition to CD19 expression, EBF1 is also known as essential gene for VDJ recombination, especially V_H_ to DJ_H_ recombination (*23*). To determine whether miR-195-transduced *Ebf1*^-/-^ cells rearranged the VDJ region, we attempt to detect V_H_-J_H_ assembled gene segments in the engrafted mouse bone marrow cells by droplet digital PCR (ddPCR). The data revealed that there were certain number of V_H_-J_H_ segments in the bone marrow of mice engrafted miR-195-transduced *Ebf1*^-/-^ cells (Fig. 2C, Fig. S1). Subsequently, to expect the EBF1 independent reconstitution enabled B cell receptor to express as IgM, we analyzed B cell populations in the engrafted mouse bone marrow. Not much but some miR-195-transduced cells expressed IgM on cell surface likely as normal immature B cell in bone marrow 10 days after engraftment (Fig. 2D). Moreover, these IgM positive cells were also detected in splenocytes. These data suggested that engrafted cells had differentiated into IgM positive immature or mature B cells, and they have been recruited to spleen. The critical function of B cell is changing B cell receptor from IgM to IgG following class switch DNA recombination, which is accompanied by stimuli-induced cell proliferation. To clarify whether miR-195-transduced *Ebf1*^-/-^ B cells have the function, whole splenocyte of the engrafted mice were stimulated with IL-4 and LPS, which causes class switch recombination to IgG1 (*24*). While control transduced GFP positive cells did not expand by the stimuli, miR-195-transduced GFP positive cells expanded enough to be surely detected and importantly a part of them expressed IgG1 (Fig. 2E). These data suggested that miR-195 has a potential to induce B cell differentiation from HPCs to mature B cells, resulting in class switch recombination even when critical regulator EBF1 is absent.

**Fig. 2.**
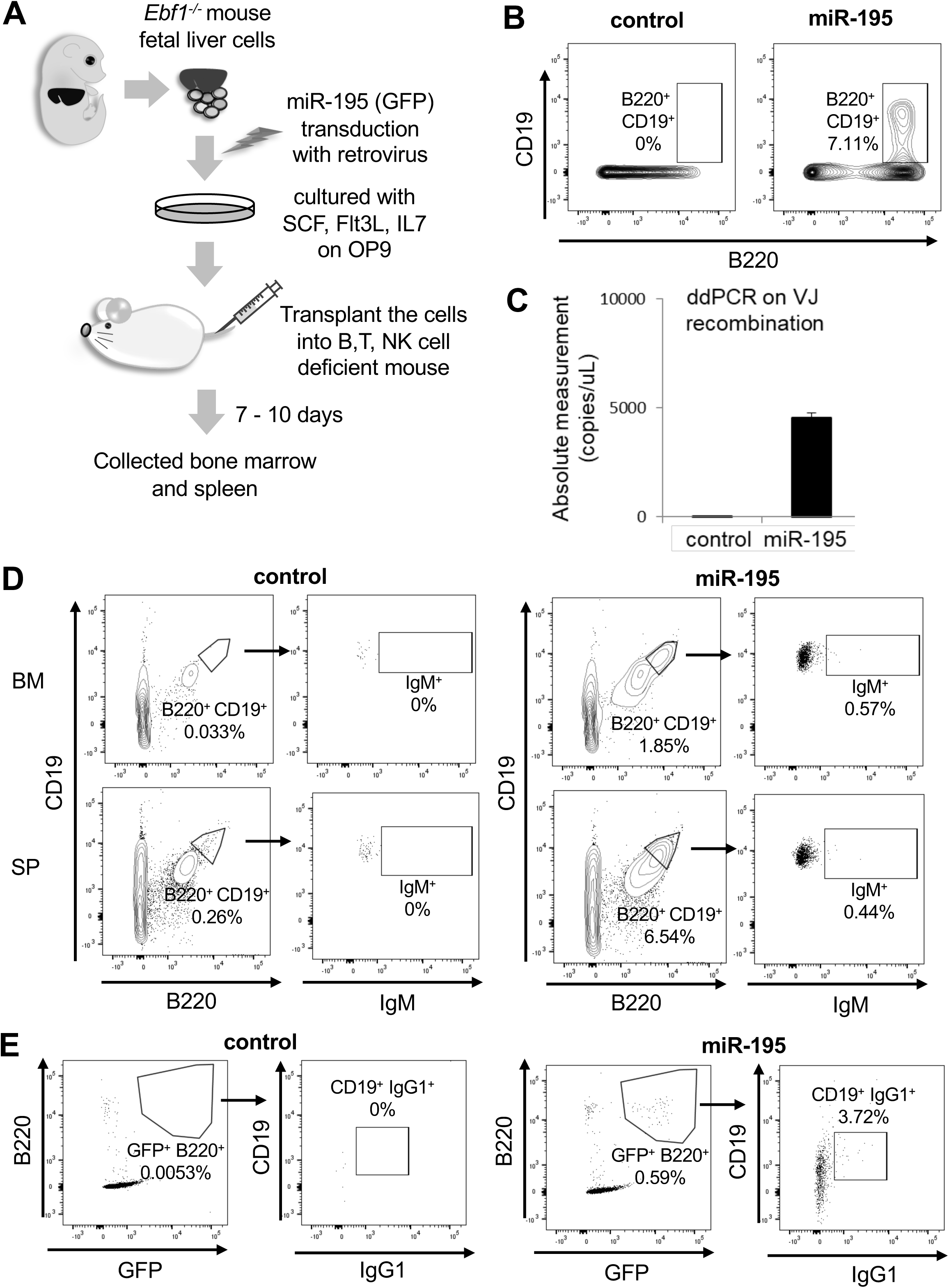
miR-195 leads *Ebf1*-deficient HPCs to mature into B cells with bone marrow niche assistance. (A) *In vivo* analysis of B cell development of *Ebf1***^-/-^** HPCs. (B) Flow cytometry analysis of control and miR-195-expressing *Ebf1***^-/-^** HPCs in the bone marrow collected at 7 days after transplantation. (C) Using droplet digital PCR, VJ region fragments were amplified from the genomic DNA of B220^+^ cells in the bone marrow of mice transplanted with control and miR-195-expressing *Ebf1***^-/-^** HPCs. (D) Flow cytometry analysis of control and miR-195-expressing *Ebf1***^-/-^**HPCs in the bone marrow (BM) and spleen (SP), at 10 days after transplantation. (E) Flow cytometry analysis of class-switch recombination. Splenocytes of mice transplanted with control and miR-195-expressing *Ebf1***^-/-^** HPCs were cultured for 72 hrs with IgG1 class-switch stimuli, LPS, and IL-4. Each flow cytometric data is representative of *n*=3.

### miR-195 physiologically maintains several B cell populations

As ectopic miR-195 expression revealed its potential in B cell development. Next, to investigate contribution of endogenous miR-195 for B cell lineage populations, miR-195 deficient mice in which the genome around miR-195-5p was eliminated by CRISPR/Cas9 system were established. The analysis of HPC lineage populations in the bone marrow revealed that several B cell-related progenitors were relatively reduced in miR-195^-/-^ mice. Sca-1^-^ c-kit^+^ common myeloid progenitor cell population was increased but controversially, Sca-1^+^ c-kit^-^ (LSK^-^) cells was decreased in miR-195^-/-^ mice (Fig. 3A). As LSK^-^ cells mainly includes early lymphoid precursor, these results suggested that miR-195 is involved in hematopoiesis including differentiation of stem cells toward lymphoid and early B cells (*25*). While analysis of each early B cell populations did not show significant difference, whole B220^+^ IgM^-^ pre B cell populations was slightly increased in the BM of miR-195^-/-^ mice (Fig. 3B). In the splenic B cells, marginal zone B (MZB) cells were reduced in miR-195^-/-^ mice (Fig. 3C). MZB cells was previously reported to be highly dependent on EBF1 activity and disappear in absence of EBF1. B-1 cells was likewise crucially regulated EBF1 as well(*26*). In the peritoneal cavity of miR-195^-/-^ mice, B-1 cells were significantly decreased (Fig. 3D). These results suggested that miR-195 contributed to maintain several EBF1-dependent mature B cell populations at least in a part. Taken together, these results were consistent with those obtained from ectopic expression of miR-195.

**Fig. 3.**
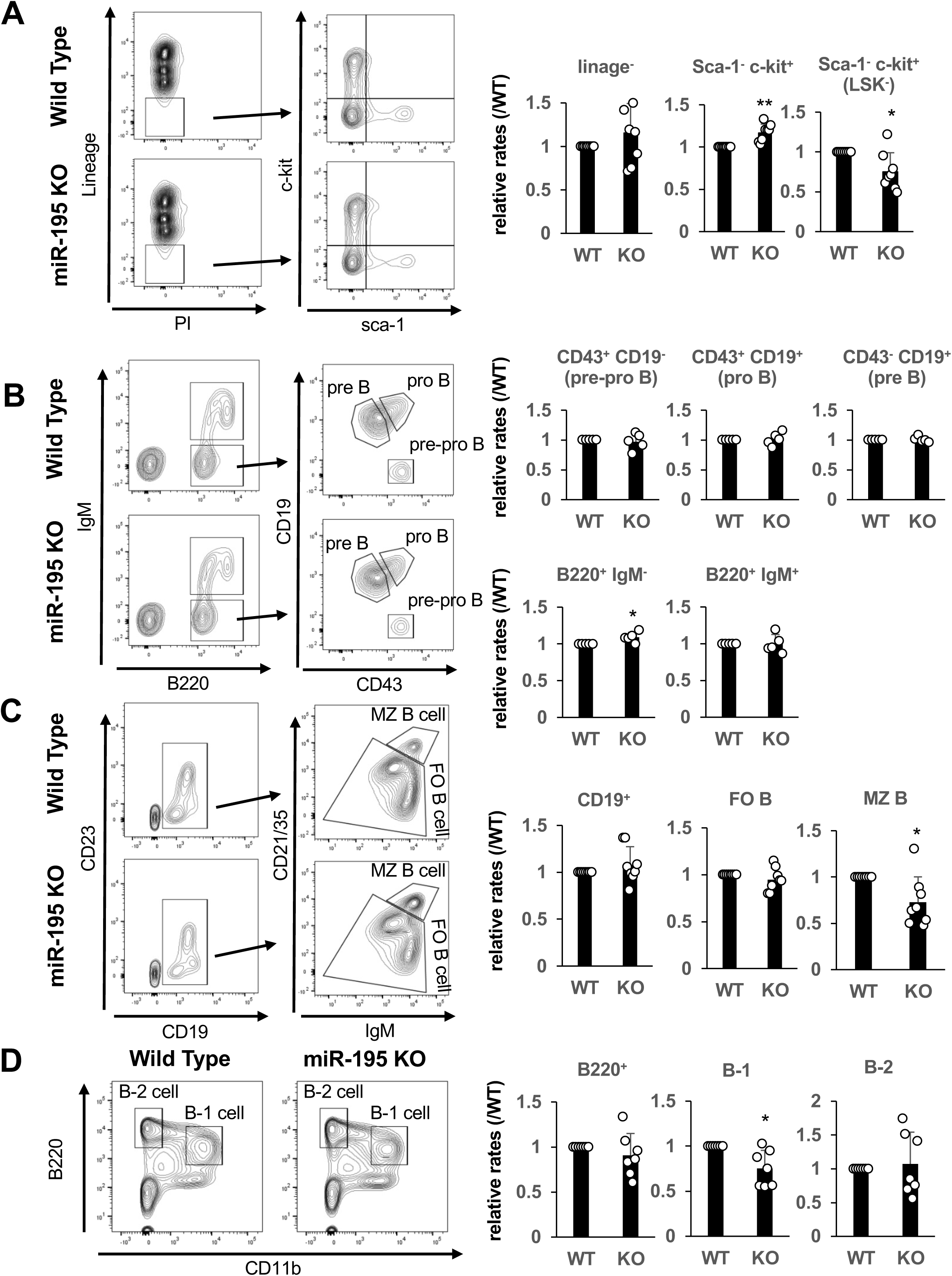
Several B cell populations are disturbed in the miR-195-deficient mice. Flow cytometry data of B cell lineage populations in miR-195**^-/-^** and littermate WT mice. Representative plots (left side) and mean ± S.D. of relative population rates in each littermate WT mice (right side) are shown. (A) Analysis of early B cell populations in the bone marrow. Pre-pro-B (B220^+^ IgM^-^ CD43^+^ CD19^-^); pro-B (B220^+^ IgM^-^ CD43^+^ CD19^+^); pre-B (B220^+^ IgM^-^ CD43^-^ CD19^+^); *n*=5. (B) Analysis of hematopoietic progenitor populations in the bone marrow; *n*=5. (C) Analysis of B cell populations in the spleen. FO B (CD19^+^ IgM^+^ CD21/35^low-middle^); MZ B (CD19^+^ IgM^+^ CD21/35^high^); *n*=8. (D) Analysis of B cell populations in the peritoneal cavity: B-1 (B220^+^ CD11b^+^); B-2 (B220^+^ CD11b^-^); *n*=7. Statistical significance was tested using one-sample *t* test. \**p*<0.05; \*\**p*<0.01. WT, wild-type.

### FOXO1 phosphorylation pathway targeted by miR-195 was responsible for B cell lineage commitment

To elucidate how miR-195 promote B cell development in EBF1 deficient HPCs, we analyzed regulatory networks of predicted miR-195 target genes by using starBase_v2.0 and David Bioinformatics Resources 6.8 in KEGG pathway database (*27–33*). Several gene regulation networks were detected as candidates of responsible pathways on the miR-195 function (Table S1 and S2). Remarkably, MAPK signaling pathway and PI3K-Akt signaling pathway included various targets of miR-195. Both MAPK and Akt were known to phosphorylate and degradate FOXO1, which was a critical factor in several stages of B cell development (*34*). Thereby, we focused on the predicted miR-195 targets: *Pik3r1*, *Pdpk1*, *Akt3*, *Raf1*, *Sos2,* and *Mapk3*, which were involved in and activate MAPK and PI3K-Akt pathways. First, to confirm that the predicted targets are actually regulated by miR-195, we picked up 3’UTR of *Mapk3* and *Akt3,* which were especially important in the pathways, and inserted in a luciferase reporter assay plasmid. As expected, the luciferase activity was down-regulated by miR-195 transduction, but it was not impaired by transduction of miR-195 mutant of mature miRNA region (Fig. 4A). Furthermore, to determine whether the predicted targets were actually regulated by miR-195, we measured the expression levels in miR-195-transduced *Ebf1*^-/-^ HPCs, and qPCR analysis showed that miR-195 transduction certainly decreased the mRNA levels (Fig. 4B).

**Fig. 4.**
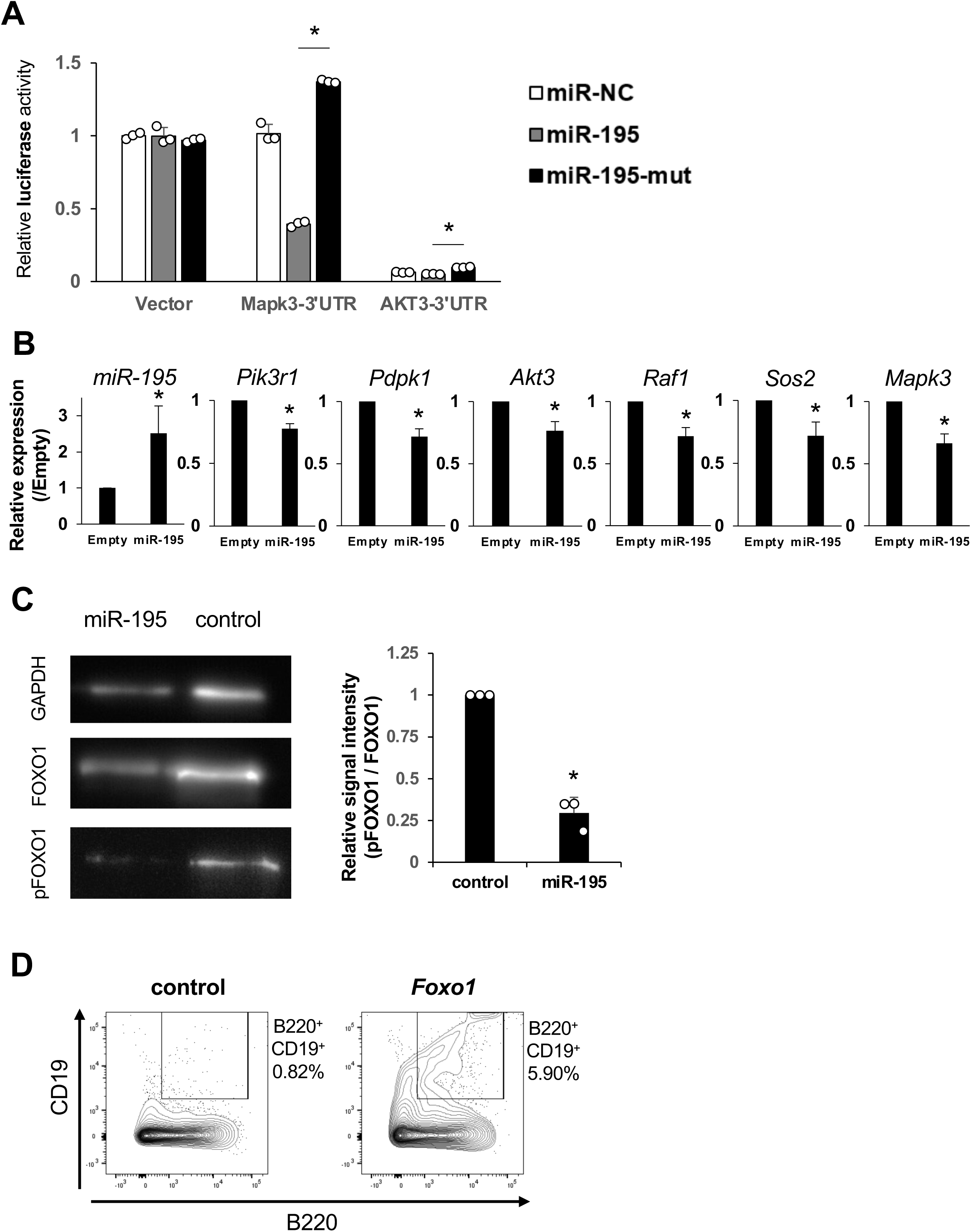
FOXO1 phosphorylation pathways are key targets of miR-195 for promotion of B cell development. (A) Relative expression rate of miR-195 and predicted target genes were compared between control (EMPTY) and miR-195-expressing *Ebf1***^-/-^** HPCs. (B) Relative luciferase inhibitory rates of miR-195 onto predicted target 3′-UTR were analyzed using Dual-Luciferase^®^ reporter assay. (C) Western blot of FOXO1 and phosphorylated FOXO1 (pFOXO1) in control and miR-195-expressing *Ebf1***^-/-^** HPCs. Quantification of FOXO1 and phospho-FOXO1 band intensities from three independent experiments is shown in the bar graph. Data are presented as mean ± SD. Shown data is representative of *n*=3. (D) Flow cytometry analysis of control and *Foxo1*-expressing *Ebf1***^-/-^** HPCs. Shown data is representative of *n*=3. Statical significance was tested using one-sample *t* test. \**p*<0.05, *n*=3.

Because of sequence similarity among miR-15/16 family members, the baseline levels detected in control samples may include signal from endogenous miRNAs such as miR-497 or miR-16. Thus, the observed increase (log_2_FC ∼2.5) may underestimate the actual level of miR-195 overexpression. In line with these findings, *Mapk3* expression was also downregulated in our microarray analysis of miR-195-transduced *Ebf1*^−/−^ cells. However, for *Akt3*, the microarray results were inconsistent across different probes, suggesting probe-dependent variability. Therefore, while qPCR and reporter assays support *Akt3* as a potential target of miR-195, its regulation remains to be further validated. Next, to evaluate inhibition of FOXO1 phosphorylation and degradation by miR-195, we compared protein levels of FOXO1 and phosphorylated FOXO1 (pFOXO1) in miR-195-transduced *Ebf1*^-/-^ HPCs. The western blotting results revealed that miR-195 transduction decreased pFOXO1 levels and increased relative FOXO1 protein levels (Fig. 4C, Fig. S2).

We also performed western blotting for PAX5 and ERG using the same samples. The results showed no significant change in these protein levels between miR-195-transduced and control Ebf1^−/−^ cells (Fig. S3), consistent with the modest upregulation observed in our microarray data. Finally, to determine whether FOXO1 accumulation is sufficient for *Ebf1*^-/-^ HPCs to differentiate into pro-B cells, *Ebf1*^-/-^ HPCs were transduced with *Foxo1* and cultured under the B cell differentiating condition. Similar to miR-195 transduction, Foxo1 transduction arose B220 and CD19 double positive *Ebf1*^-/-^ cells, which was accompanied with CD19 positive but B220 negative population (Fig. 4D). These data indicated that FOXO1 accumulation by inhibition of phosphorylating pathways was responsible for *Ebf1*^-/-^ HPCs to differentiate into B cell lineage.

### Epigenetically activated genes in pro-B cells by miR-195 is fewer than by EBF1

In B cell development, epigenetic changes of transcription factors and differentiation molecules are crucial for proper development, which are mainly regulated by EBF1 (*35*, *36*). We investigated transposase-accessible chromatin using deposited sequencing data (ATAC-seq) of *Ebf1*^-/-^ pro-B cells and wild type pro-B cells from GSE92434 and cells in early B cell lineages from GSE100738. While wild type pro-B cells/ *Ebf1*^-/-^ pro-B cells differentially accessible (DA) ATAC peaks were observed in 2809 sites, wild type CD19 positive/ CD19 negative early B cells DA ATAC peaks were in 904 sites. Then, 678 sites were overlapped, which were considered to be regulated by EBF1 as important locus for early B cell development. Moreover, some of them were overlapped with miR-195-transduced B220 and CD19 double positive *Ebf1*^-/-^ cells (miR-195 CD19+) / B220 positive CD19 negative *Ebf1*^-/-^ cells (control CD19-) DA ATAC peaks (73 out of 226 peaks), which were considered to be regulated by miR-195 (Fig. 5B). These peaks included important genes for early B cell development, such as *Pax5*, *Runx1*, *Erg*, *Ifr8*, and *Blnk*, and B cell-related genes, such as *Rarres1*, *Ciita*, and *Atg7* (Table S3). These results indicated that gene locus opened by miR-195 were fewer than by EBF1, but they included several key locus for B cell differentiation, and they were enough to differentiate the progenitor cells to mature B cells. Moreover, HOMER Motif Analysis revealed that enriched motives opened by EBF1 and by miR-195 were 198 and 111, respectively (Fig. 5C). The common motives were 104 which included critical genes for B cell development, such as E2A, Foxo1, and PAX5, and high ranked motives were very similar between EBF1 and miR-195 (Fig. 5D). These results suggested that miR-195 transduction opened important chromatin regions for early B cells, which were normally regulated by EBF1.

**Fig. 5.**
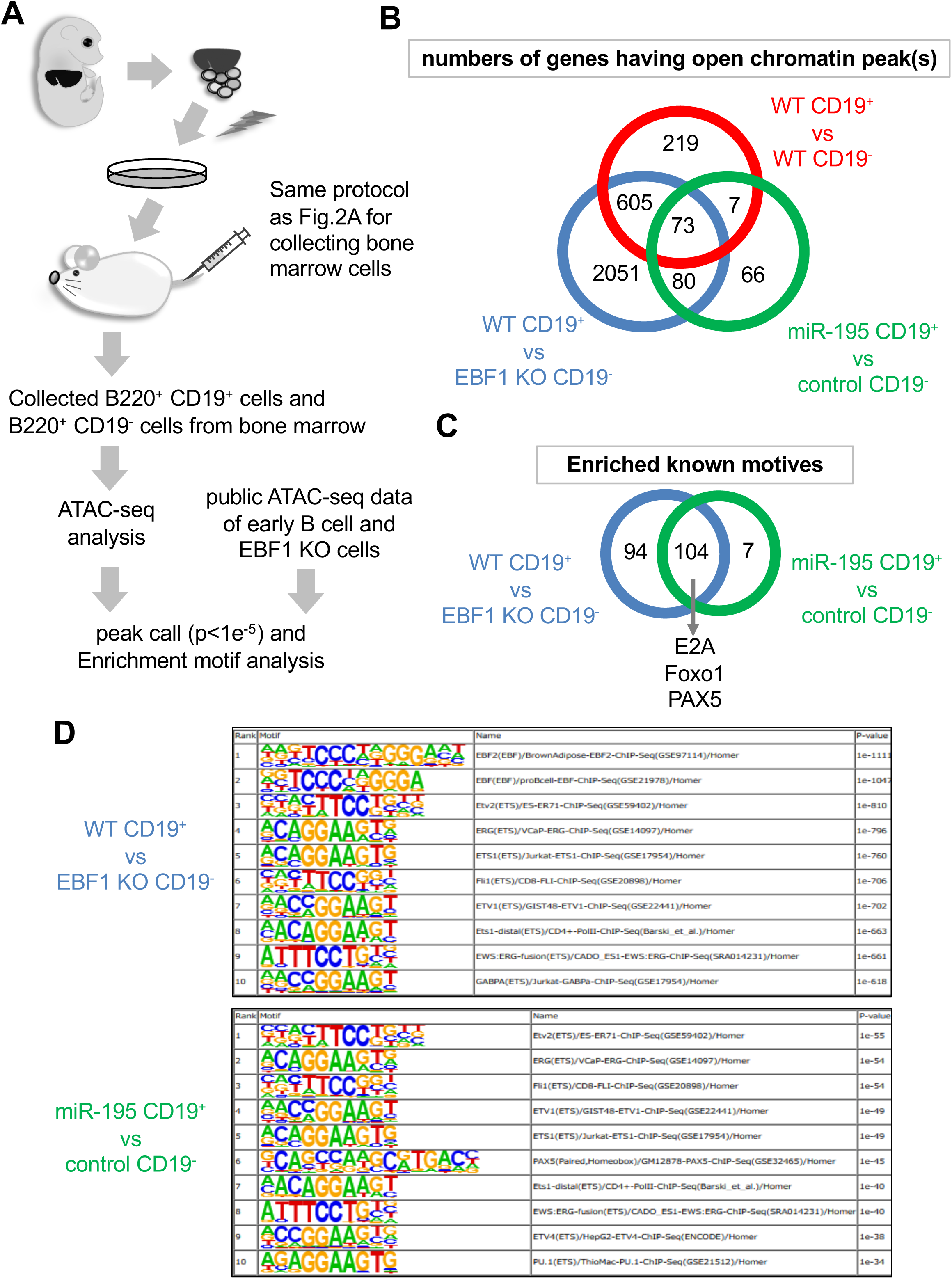
ATAC-seq analysis of *Ebf1*^-/-^ CD19-positive B cells differentiated by miR-195. (A) Outline of analysis of open chromatin regions in miR-195-expressing *Ebf1***^-/-^** cells. (B) Venn diagram of numbers of genes in which DNA regions of open chromatin peaks were detected by means of peak call analysis. The analyses were examined between CD19-negative (FrA) and-positive (FrB, FrC, and FrD) stages of B cell development (GSE100738; upper red circle); wild-type and *Ebf1***^-/-^** pro-B cells (GSE92434; left-lower blue circle); B220^+^ CD19^-^ cells of control and B220^+^ CD19^+^ positive miR-195-expressing *Ebf1***^-/-^** cells (right-lower green circle). Overlapping regions in the Venn diagram are interpreted as follows: the intersection of WT and Rescue represents canonical EBF1-regulated regions; the overlap between Rescue and miR-195 indicates partial mimicry by miR-195; and regions unique to miR-195 may reflect EBF1-independent chromatin changes. (C and D) Venn diagram of numbers of enriched known motifs detected using HOMER find motif analysis (C) and lists of high *p*-value motifs, up to rank 10 (D).

Finally, we concluded that miR-195 transduction was able to compensate EBF1 deficiency in B cell development through activation of FOXO1 and epigenetic regulation of several B cell-related genes.

## Discussion

The canonical notion of hematopoietic fate determination implies that EBF1 is an indispensable factor for B lymphopoiesis. However, in this study, we showed that a single microRNA miR-195 rescued the arrest of pro-B cell differentiation induced by EBF1 deficiency. As miRNA plays roles in a bundle of their family, single miRNA deficient mice often do not show significant phenotype (*37*). Nevertheless, miR-195 deficient mice showed not much but sure decreased number of several hematopoietic cells including marginal zone B cells and peritoneal B-1 cells, which were reported to almost disappear in EBF1^ihCd2^ mice in which EBF1 was deficient in mature B cells (*38*). Considering that other miRNA deficient mice have subtle phenotype and miR-195 is one of the large family, including miR-15/16 and miR-195/497 (*39*), the remarkable potential of miR-195 is beyond a fine tuner as microRNA, at least as far as it is considered with regard to B cell lineage commitment.

A part of the mechanisms of the potent function of miR-195 was caused by inhibition of phosphorylation of FOXO1. FOXO1 is a transcriptional factor controlled by EBF1 and strongly promote differentiation of pre-B cell. FOXO1 activity is regulated by PI3K/AKT pathway and several miRNAs were reported to be involved in the regulation (*40*). We showed *Foxo1* transduction enable EBF1 deficient cell to express CD19.

However, the CD19 positive cells rapidly disappeared and couldn’t be detected in transplanted mice (data not shown). It is presumable that FOXO1 activity was necessary to express CD19, but other factors undertake maintenance and proliferation of the developing cells. ATAC-seq analysis revealed that miR-195 was directly or indirectly involved in chromatin accessibility. As the chromatin regions and motives opened by miR-195 were critical for B cell differentiation and hematopoiesis, further investigation is needed for the mechanism.

Although our study indicates that miR-195 has the potential to promote B cell lineage commitment in the absence of EBF1, the precise downstream targets and mechanisms remain only partially defined. We hypothesize that the observed effects are mediated through the downregulation of multiple mRNA targets involved in opposing B-lineage differentiation, including kinases in the MAPK and PI3K-Akt pathways that modulate FOXO1 phosphorylation. While our microarray, qPCR, and luciferase assays support the regulation of specific targets such as *Mapk3* and *Akt3*, a more comprehensive identification of direct targets—especially those related to transcriptional and epigenetic regulation—would further strengthen our conclusions. We interpret our findings as revealing the potential of miR-195 to compensate for EBF1 deficiency, rather than a demonstration of its physiological role. Future studies using global transcriptome, proteome, and chromatin-binding assays will be essential to fully elucidate the mechanisms underlying this observation.

To compensate for the lack of transcriptome data from sorted *miR-195*-transduced pre-pro-B or CD19+ *Ebf1*^−/−^ cells, we compared our microarray data with publicly available RNA-seq profiles of *Ebf1*^−/−^ pro-B cells (GSE92434). This analysis revealed that several B-lineage defining genes downregulated in *Ebf1* deficiency were upregulated upon miR-195 expression, suggesting that miR-195 may partially restore transcriptional programs disrupted by the loss of EBF1.

Although direct evidence of FOXO1 binding to B-lineage gene loci (e.g., via ChIP-seq or CUT&RUN) is currently lacking due to technical limitations in cell numbers, our results suggest that FOXO1 plays a key functional role. This is supported by its increased protein level upon miR-195 expression, the partial phenocopy by FOXO1 overexpression, and the enrichment of FOXO1 motifs in open chromatin regions identified by ATAC-seq. Future studies incorporating FOXO1 chromatin profiling will be important to validate its direct regulatory role in this context.

While ddPCR provided a sensitive means to detect *VH-JH* rearranged fragments, it does not offer resolution of specific V, D, or J gene usage or recombination completeness.

Therefore, the full extent and diversity of V(D)J recombination in *Ebf1*^−/−^ miR-195-induced CD19⁺ cells remains to be clarified. Future studies incorporating high-throughput sequencing approaches will be important to fully characterize the immunoglobulin repertoire and confirm progression through the pre-BCR checkpoint.

While our data support the B-lineage identity of miR-195-induced *Ebf1*^−/−^ CD19⁺ cells based on gene expression, chromatin accessibility, and immunoglobulin expression, we have not directly tested their lineage plasticity under alternative differentiation conditions. Whether these cells retain responsiveness to myeloid cytokines or exhibit residual multipotency remains to be determined. Future studies using single-cell fate mapping or in vitro differentiation assays will be required to fully define the lineage commitment status of this population.

While our results demonstrate that ectopic expression of miR-195 can compensate for the loss of EBF1 in promoting B cell development, we acknowledge that this does not necessarily reflect a physiological role for miR-195. The miR-195 knockout mice exhibited only mild alterations in B cell populations, suggesting that under normal conditions, miR-195 is not essential for B lymphopoiesis. Therefore, our findings should be interpreted as highlighting the potential of miR-195 to modulate B cell fate under specific conditions, rather than indicating its requirement in physiological B cell development. Further studies will be needed to determine whether miR-195 plays a more prominent role under stress or disease contexts, or in cooperation with other miRNAs.

The luciferase activity was markedly reduced in the presence of the Akt3 3’UTR, even in cells transduced with a control vector (Fig. 4A). We hypothesize that the Akt3 3’UTR contains strong post-transcriptional regulatory elements—such as AU-rich elements or binding sites for endogenous miRNAs or RNA-binding proteins—which may suppress mRNA stability or translation independent of miR-195. Alternatively, the secondary structure or length of the UTR may inherently reduce luciferase expression.

## Materials and Methods

### Plasmid construction

To construct MDH1-PGK-GFP-miR-195, genomic DNA was first extracted from RS4;11 using the DNeasy Tissue Extraction Kit (Qiagen). Next, a segment around miR-195 was amplified by means of PCR, using Pfx polymerase (Invitrogen) and the oligonucleotides, 5′-AGATCTCTCGAGAAGGAGAGGGTGGGGTAT-3′ and 5′ - GGGGCGGAATTCGCTATTCCCGCATAAGCA-3′. The obtained PCR product was then cloned into the XhoI-EcoRI site of MDH1-PGK-GFP 2.0 (Addgene #11375). To construct pMYs-RFP-Foxo1, first, pEX-Foxo1 (in which mouse Foxo1 is optimized for gene synthesis; Eurofins Genomics K.K.) was synthesized and inserted into the EcoRI-XhoI site of pEX. Next, the Foxo1 region was extracted using the restriction enzymes and inserted into pMYs-RFP retroviral vector (kindly provided by Prof. T. Kitamura, Tokyo University). For in vitro transcription of small-guide RNA (sgRNA), pUC57-195sg-upstream and-downstream were generated. Both plasmids originated from the pUC57-sgRNA expression vector (Addgene #51132), and the annealed oligonucleotides were inserted into a BsaI site (For the former, 5′ - TAGGCCCACAAAGGCAGGGACCTA-3′ and 5′ - AAACTAGGTCCCTGCCTTTGTGGG-3′ were annealed, while for the latter, 5′ - TAGGGGAAGTGAGTCTGCCAATAT-3′ and 5′ - AAACATATTGGCAGACTCACTTCC-3′ were annealed). For the Dual-Luciferase® assay, psiCHECK-2 vector was purchased from Promega and the 3′-UTRs of Akt3 and Mapk3 were inserted between the XhoI and NotI sites. MDH1-PGK-GFP-miR-195-mut was generated by mutating 6 bases, from the second to seventh bases of the mature miR-195 and complimentary regions of the stem loop structure in MDH1-PGK-GFP-miR-195. In detail, normal miR-195 stem loop sequence 5′ - AGCUUCCCUGGCUCUAGCAGCACAGAAAUAUUGGCACAGGGAAGCGAGUCUGCCAAUAUUGGCUGUGCUGCUCCAGGCAGGGUGGUG-3′ (mature miR-195-5p sequence 5′-UAGCAGCACAGAAAUAUUGGC-3′) was mutated to 5′ - AGCUUCCCUGGCUCUgcgccgACAGAAAUAUUGGCACAGGGAAGCGAGUCUGCCAAUAUUGGCUGUcggcgcCCAGGCAGGGUGGUG-3′ (mature sequence 5′ - UgcgccgACAGAAAUAUUGGC-3′).

### Animals

C57BL/6 mice were purchased from CLEA Japan Inc. NOD/Shi-scid, IL-2RγKO (NOG) and B6RG mice were purchased from Central Institute for Experimental Animals (CIEA). The Ebf1^−/+^ mice were originally generated by R. Grosschedl (*41*). miR-195-deficient mice were generated based on the CRISPR/Cas9 system established by C. Gurumurthy (*42*), using pUC57-195sg-upstream and-downstream for sgRNA expression and pBGK (Addgene #65796) for Cas9 mRNA expression. Sanger sequencing confirmed a deletion of 5,103 base pairs at chromosome 11 (GRCm38/mm10 chr11:70,234,425–70,235,103), encompassing the entire *miR-497* sequence upstream and 61 bp of the 93-bp *miR-195* precursor. The deletion was validated using genomic DNA and aligned to the mouse reference genome. All transgenic mice used for experiments were backcrossed to the C57BL/6 background for at least eight generations to minimize off-target effects. The obtained mice were subsequently bred and housed at Tokai University. All the animal experiments in this study complied with the Guidelines for the Care and Use of Animals for Scientific Purposes at Tokai University. To reduce the number of sacrificed animals, the sample sizes for each animal experiment were empirically determined from previous studies or the results of the first littermate mice.

### Flow cytometry analysis

Cells were collected and washed in FACS buffer (phosphate-buffered saline supplemented with 2% fetal bovine serum) and subsequently stained with the following antibodies purchased from BioLegend: anti-c-kit (2B8),-Sca-1 (D7),-IL7Rα (A7R34),-B220 (RA3-6B2),-IgM (RMM-1),-CD3ε (145-2C11),-CD4 (GK1.5),-CD8 (53-6.7),-CD11b (M1/70),-CD19 (1D3),-CD23 (B3B4), and-IgG1 (RMG1-1) and Thermo Fisher: anti-Flt3 (A2F10),-CD43 (eBioR2/60), and-CD21/35 (eBio8D9). All samples were analyzed on the BD FACSVerse™ system and the data obtained was analyzed using FlowJo. FACSAria™ III was used for cell sorting.

### Culture of lineage-negative (Lin^-^) cells from the fetal liver

Fetal livers were harvested from pregnant C57BL/6 or *Ebf1^+/−^*mice (mated with *Ebf1^+/−^*male) at 13.5 days after vaginal plug formation and minced gently by means of pipetting. The cell suspensions were filtered through a 67-µm pore nylon mesh and Lin^-^ cells were collected using the Lineage Cell Depletion Kit, mouse and AutoMACS^®^ Pro Separator (Miltenyi Biotec), according to the manufacturer’s instructions. Subsequently, the collected Lin^-^ cells were transduced with miR-195 or *Foxo1* by means of retroviral transfection. In brief, Platinum-E cells were transfected with MDH1-PGK-GFP (for EMPTY sample) or MDH1-PGK-GFP-miR-195 or pMYs-RFP-Foxo1 using PEI MAX^®^ (Polysciences Inc.), and retroviral supernatants were harvested 48 hours later. Lin^-^ cells were infected with the supernatants using 10 µg/mL Polybrene (Sigma-Aldrich). The infected and transduced Lin^-^ cells were cultured and differentiated into B cells on OP9 cells in IMDM (Thermo Fisher) supplemented with 10% fetal bovine serum, 1 mM sodium pyruvate, 0.1 mM non-essential amino acid solution, 50 µM 2-mercaptoethanol, 100 units/mL penicillin G, 100 µg/mL streptomycin (all from Wako), and 10 ng/mL recombinant SCF, IL-7, and Flt3-ligand (R&D Systems). Cells were cultured on OP9 cells for 7 days before analysis unless otherwise specified. For *in vivo* analysis of B cell development of EBF1^-/-^ Lin^-^ cells, 1×10^6^ cells were injected into the NOG or B6RG mice after >7 days of culture and expansion *in vitro*.

### Microarray analysis

Total RNA was isolated using the RNeasy MINI Kit (Qiagen), and its quality was analyzed using the 2100 Bioanalyzer (Agilent Technologies). Approximately 100 ng RNA was labeled, and gene expression microarray analysis was performed using the Agilent Whole Mouse Genome Microarray 4×44K v2 (Agilent Technologies), according to the manufacturer’s instructions. The processed data was analyzed using GeneSpring GX version 14.9 (Agilent Technologies). Raw intensity values were normalized using the 75^th^ percentile and transformed to the Log_2_ scale. All experiments were carried out in duplicates.

### Droplet digital PCR (ddPCR)

To carry out ddPCR for VJ recombination analysis, total DNA was isolated from whole cells of the bone marrow in miR-195-transduced *Ebf1*^-/-^ FL HPCs-engrafted NOG mice, using the Wizard^®^ Genomic DNA Purification Kit (Promega). ddPCR was conducted using QX100 Droplet Digital PCR system (Bio-Rad). Briefly, 3.3 μL of template cDNA with 20× primer and a TaqMan™ probe set was partitioned into approximately 20,000 droplets using the QX100 Droplet Generator, for amplification. The cycling conditions were 95°C for 10 min, followed by 50 cycles of 95°C for 15 sec and 60°C for 1 min, and a final 10-min incubation at 98°C. The droplets were subsequently read automatically using the QX10 droplet reader. The data were analyzed with QuantaSoft analysis software (ver. 1.3.2.0; Bio-Rad). The primers used were as follows: forward primer – 5′-GAGGACTCTGCRGTCTATTWCTGTGC-3′; reverse primer – 5′-CCCTGACCCAGACCCATGT-3′; and probe – 5′-6FAM-TTCAACCCCTTTGTCCCAAAGTT-TAM-3′.

### Class-switch stimulation

EBF1^-/-^ Lin^-^ cells were transduced with EMPTY and miR-195-expressing vector and transplanted into B6RG mice. At 10 days post-transplantation, the spleens were collected from the mice, minced with slide glasses, and filtered through a 67-µm pore nylon mesh. IgM^+^ cells were sorted and stimulated for 3 days with 12.5 μg/mL lipopolysaccharide (Sigma-Aldrich) and 7.5 ng/mL IL-4 (Peprotech) in RPMI-1640 (Wako) supplemented with 10% fetal bovine serum, 100 U/mL penicillin G, and 100 µg/mL streptomycin.

### Gene Ontology analysis

The miR-195 targetomes were gathered from the miR-195 target mRNAs identified from three databases (Targetscan, miRDB, and microRNA.org) and by comparing the microarray data of the targets in control-and miR-195-transduced Ebf1^-/-^ FL HPCs. To investigate the biological functions, these genes were applied to the Gene Ontology classification using GeneSpringGX11.

### Quantitative real-time PCR

For mRNA quantification, total RNA was isolated using Sepasol-RNA I Super G (Nacalai Tesque) and cDNA was synthesized from it using the ReverTra Ace™ qPCR RT Master Mix (TOYOBO). qPCR was performed using THUNDERBIRD™ SYBR^®^ qPCR Mix (TOYOBO) on the StepOnePlus™ Real-Time PCR System (Thermo Fisher). The following primers were used for qPCR: Pik3r1 – 5′-AAACTCCGAGACACTGCTGA-3′ and 5′-GAGTGTAATCGCCGTGCATT-3′; Pdpk1 – 5′-CTGGGCTCTGCTCTAGTGTT-3′ and 5′-CCCAGGTTCAGGACAGGATT-3′; Akt3 – 5′-GTGGACCACTGTTATAGAGAGAACAT-3′ and 5′-TTGGATAGCTTCCGTCCACT-3′; Raf1 – 5′-TCTTCCATCGAGCTGCTTCA-3′ and 5′-GGATGTAGTCAGCGTGCAAG-3′; Sos2 – 5′-AACTTTGAAGAACGGGTGGC-3′ and 5′-TTTCCTGCAGTGCCTCAAAC-3′; and Mapk3 – 5′-ACTACCTGGACCAGCTCAAC-3′ and 5′-TAGGAAAGAGCTTGGCCCAA-3′. For miR-195 quantification, TaqMan™ MicroRNA Assay (ABI) was used. Briefly, total RNA was isolated using Sepasol-RNA I Super G and cDNA was synthesized from it using the microRNA TaqMan™ MicroRNA Reverse Transcription Kit (Thermo Fisher) and a specific primer, 5′-UAGCAGCACAGAAAUAUUGGC-3′. The expression levels were measured using the TaqMan™ Fast Advanced Master Mix (Thermo Fisher) on the StepOnePlus™ Real-Time PCR System. Given the high sequence similarity among miR-15/16 family members, the TaqMan assay for miR-195 may detect related miRNAs such as miR-16. Therefore, we interpreted miR-195 qPCR results as approximate estimates rather than precise quantification. GAPDH was used for normalization to maintain consistency with other qPCR assays in this study. All reagents and kits in this section were used according to the manufacturer’s instructions. Target RNA expression levels were compared with those of GAPDH using the 2^-ΔΔCt^ method.

### Dual-Luciferase**^®^** assay

293T cells were co-transfected with 20 ng psiCHECK-2 of *Akt3* or *Mapk3* and 100 ng MDH1-PGK-GFP-miR-195 or MDH1-PGK-GFP-miR-195-mut. At 48 hrs post-transfection, the relative amounts of Renilla and firefly luciferase were analyzed using a Dual-Luciferase^®^ Reporter Assay System (Promega). The Renilla/firefly luciferase ratio was calculated and normalized against the control.

### Western blot

Total proteins were collected from whole cells using radioimmunoprecipitation assay buffer (Wako) with protease inhibitor cocktail (Sigma-Aldrich) and SDS sample buffer (60 mM Tris-HCl pH 6.8, 2% SDS, 10% glycerol, and 50 mM dithiothreitol). The proteins were separated using SDS-PAGE and the western blot signal was detected and analyzed using the Immobilon Western Chemiluminescent HRP Substrate (Millipore) on Ez-Capture MG AE-9300 (ATTO). The following antibodies were used: anti-FOXO1 (C29H4, Cell Signaling Technology),-phospho-FOXO1(Ser256) (9461, Cell Signaling Technology), and-GAPDH (G9545, Sigma-Aldrich). Signal intensities were quantified using ImageJ version 1.54g (*43*).

### ATAC-seq analysis

For ATAC-seq analysis, B220^+^ CD19^+^ and B220^+^ cells were sorted from the bone marrow of NOG mice transplanted with miR-195-transduced EBF1^-/-^ Lin^-^ cells. B220^+^ cells were also sorted from the empty transduced sample. The collected cells were resolved using CELLBANKER^®^ (Takara Bio) and temporarily preserved at –20°C.

ATAC-seq libraries were prepared from the cryopreserved cells according to the Omni-ATAC protocol (*44*). Briefly, >5,000 cells were lysed and subjected to a transposition reaction. The transposed fragments were pre-amplified, quantitated using RT-PCR, and then amplified again. The prepared libraries were sequenced on the NextSeq 550 platform (Illumina) with paired-end reads (read 1, 75 bp; index 1, 8 bp; index 2, 8 bp; read 2, 75 bp). Short-read data were trimmed using sickle 1.33 (https://github.com/najoshi/sickle) and mapped onto a mm10 reference genome using bowtie2. Unmapped, multi, chrM mapping, and duplicate reads were eliminated using samtools 1.16.1 and Picard Tools (Picard MarkDuplicates; http://broadinstitute.github.io/picard). Peak summits in all populations were determined using the MACS3 functions (-callpeak-p 1e-5 https://github.com/macs3-project/MACS). Motif enrichment analysis was carried out using HOMER, with default settings.

## Statistical analysis

One-sample *t*-test was used to analyze differences between groups, and *p*-values<0.05 were considered statistically significant. All analyses were performed using Excel (Microsoft). Statistical significance was determined using the Fisher’s exact test, followed by multiple test corrections using the Benjamini and Yekutieli false discovery rate method.

## Acknowledgments

We thank N. Kurosaki, K. Takahashi, E. Nagashima, and members of the Department of Innovative Medical Science at Tokai University for their assistance, advice, and helpful discussions. We also thank the Support Center for Medical Research and Education at Tokai University for their technical assistance.

## Author contributions

A.K. designed the research; Y.M., T.K., R.Y., R.K., K.K., R.-K.N., and K.O. performed the research; T.I., K. Hirano, H.H., K. Hozumi, M. Ohtsuka, T. Kishikawa, C.S., M. Otsuka, R.M., K.A., and T. Kurosaki contributed new reagents and analytic tools; H.K. analyzed the data; Y.M., T.K., and A.K. wrote the paper.

## Data and materials availability

The microarray data was deposited in Gene Expression Omnibus with the identifier GSE246669, and the ATAC-seq data was also deposited with the identifier GSE246530. The other data generated in this study are available in the manuscript or supplementary materials.

## References

1. J. Zhu, S. G. Emerson, Hematopoietic cytokines, transcription factors and lineage commitment. Oncogene 21, 3295–3313 (2002).

2. T. Ikawa, H. Kawamoto, L. Y. T. Wright, C. Murre, Long-Term Cultured E2A-Deficient Hematopoietic Progenitor Cells Are Pluripotent. Immunity 20, 349–360 (2004).

3. Y. C. Lin, S. Jhunjhunwala, C. Benner, S. Heinz, E. Welinder, R. Mansson, M. Sigvardsson, J. Hagman, C. A. Espinoza, J. Dutkowski, T. Ideker, C. K. Glass, C. Murre, A global network of transcription factors, involving E2A, EBF1 and Foxo1, that orchestrates B cell fate. Nature Immunology 11, 635–643 (2010).

4. J. M. R. Pongubala, D. L. Northrup, D. W. Lancki, K. L. Medina, T. Treiber, E. Bertolino, M. Thomas, R. Grosschedl, D. Allman, H. Singh, Transcription factor EBF restricts alternative lineage options and promotes B cell fate commitment independently of Pax5. Nature Immunology 9, 203–215 (2008).

5. I. Györy, S. Boller, R. Nechanitzky, E. Mandel, S. Pott, E. Liu, R. Grosschedl, Transcription factor Ebf1 regulates differentiation stage-specific signaling, proliferation, and survival of B cells. Genes Dev. 26, 668–682 (2012).

6. K. L. Medina, J. M. R. Pongubala, K. L. Reddy, D. W. Lancki, R. DeKoter, M. Kieslinger, R. Grosschedl, H. Singh, Assembling a Gene Regulatory Network for Specification of the B Cell Fate. Developmental Cell 7, 607–617 (2004).

7. D. P. Bartel, MicroRNAs: Target Recognition and Regulatory Functions. Cell 136, 215–233 (2009).

8. R. W. Carthew, E. J. Sontheimer, Origins and Mechanisms of miRNAs and siRNAs. Cell 136, 642–655 (2009).

9. D. Cifuentes, H. Xue, D. W. Taylor, H. Patnode, Y. Mishima, S. Cheloufi, E. Ma, S. Mane, G. J. Hannon, N. D. Lawson, S. A. Wolfe, A. J. Giraldez, A Novel miRNA Processing Pathway Independent of Dicer Requires Argonaute2 Catalytic Activity. Science 328, 1694–1698 (2010).

10. N. Yamakawa, K. Okuyama, J. Ogata, A. Kanai, A. Helwak, M. Takamatsu, K. Imadome, K. Takakura, B. Chanda, N. Kurosaki, H. Yamamoto, K. Ando, H. Matsui, T. Inaba, A. Kotani, Novel functional small RNAs are selectively loaded onto mammalian Ago1. Nucleic Acids Res 42, 5289–5301 (2014).

11. C. Sevignani, G. A. Calin, L. D. Siracusa, C. M. Croce, Mammalian microRNAs: a small world for fine-tuning gene expression. Mamm Genome 17, 189–202 (2006).

12. M. Listowski, E. Heger, D. Bogusławska, B. Machnicka, K. Kuliczkowski, J. Leluk, A. Sikorski, microRNAs: fine tuning of erythropoiesis. Cellular and Molecular Biology Letters 18, 34–46 (2013).

13. S. Monticelli, K. M. Ansel, C. Xiao, N. D. Socci, A. M. Krichevsky, T.-H. Thai, N. Rajewsky, D. S. Marks, C. Sander, K. Rajewsky, A. Rao, K. S. Kosik, MicroRNA profiling of the murine hematopoietic system. Genome Biology 6, R71 (2005).

14. J. R. Neilson, G. X. Y. Zheng, C. B. Burge, P. A. Sharp, Dynamic regulation of miRNA expression in ordered stages of cellular development. Genes Dev. 21, 578– 589 (2007).

15. C.-Z. Chen, L. Li, H. F. Lodish, D. P. Bartel, MicroRNAs Modulate Hematopoietic Lineage Differentiation. Science 303, 83–86 (2004).

16. S. B. Koralov, S. A. Muljo, G. R. Galler, A. Krek, T. Chakraborty, C. Kanellopoulou, K. Jensen, B. S. Cobb, M. Merkenschlager, N. Rajewsky, K. Rajewsky, Dicer Ablation Affects Antibody Diversity and Cell Survival in the B Lymphocyte Lineage. Cell 132, 860–874 (2008).

17. C. Xiao, D. P. Calado, G. Galler, T.-H. Thai, H. C. Patterson, J. Wang, N. Rajewsky, T. P. Bender, K. Rajewsky, MiR-150 Controls B Cell Differentiation by Targeting the Transcription Factor c-Myb. Cell 131, 146–159 (2007).

18. K. Okuyama, T. Ikawa, B. Gentner, K. Hozumi, R. Harnprasopwat, J. Lu, R. Yamashita, D. Ha, T. Toyoshima, B. Chanda, T. Kawamata, K. Yokoyama, S. Wang, K. Ando, H. F. Lodish, A. Tojo, H. Kawamoto, A. Kotani, MicroRNA-126–mediated control of cell fate in B-cell myeloid progenitors as a potential alternative to transcriptional factors. PNAS 110, 13410–13415 (2013).

19. H. Qiu, J. Zhong, L. Luo, Z. Tang, N. Liu, K. Kang, L. Li, D. Gou, Regulatory Axis of miR-195/497 and HMGA1-Id3 Governs Muscle Cell Proliferation and Differentiation. International Journal of Biological Sciences 13, 157–166 (2017).

20. A. Dueñas, A. Expósito, M. del M. Muñoz, M. J. de Manuel, A. Cámara-Morales, F. Serrano-Osorio, C. García-Padilla, F. Hernández-Torres, J. N. Domínguez, A. Aránega, D. Franco, MiR-195 enhances cardiomyogenic differentiation of the proepicardium/septum transversum by Smurf1 and Foxp1 modulation. Sci Rep 10, 9334 (2020).

21. A. C. Gomes, T. Hara, V. Y. Lim, D. Herndler-Brandstetter, E. Nevius, T. Sugiyama, S. Tani-ichi, S. Schlenner, E. Richie, H.-R. Rodewald, R. A. Flavell, T. Nagasawa, K. Ikuta, J. P. Pereira, Hematopoietic Stem Cell Niches Produce Lineage-Instructive Signals to Control Multipotent Progenitor Differentiation. Immunity 45, 1219–1231 (2016).

22. K. Tokoyoda, T. Egawa, T. Sugiyama, B.-I. Choi, T. Nagasawa, Cellular Niches Controlling B Lymphocyte Behavior within Bone Marrow during Development. Immunity 20, 707–718 (2004).

23. J. M. R. Pongubala, D. L. Northrup, D. W. Lancki, K. L. Medina, T. Treiber, E. Bertolino, M. Thomas, R. Grosschedl, D. Allman, H. Singh, Transcription factor EBF restricts alternative lineage options and promotes B cell fate commitment independently of Pax5. Nat Immunol 9, 203–215 (2008).

24. M. Muramatsu, V. S. Sankaranand, S. Anant, M. Sugai, K. Kinoshita, N. O. Davidson, T. Honjo, Specific Expression of Activation-induced Cytidine Deaminase (AID), a Novel Member of the RNA-editing Deaminase Family in Germinal Center B Cells*. Journal of Biological Chemistry 274, 18470–18476 (1999).

25. R. Kumar, V. Fossati, M. Israel, H.-W. Snoeck, Lin−Sca1+Kit− Bone Marrow Cells Contain Early Lymphoid-Committed Precursors That Are Distinct from Common Lymphoid Progenitors. The Journal of Immunology 181, 7507–7513 (2008).

26. B. Vilagos, M. Hoffmann, A. Souabni, Q. Sun, B. Werner, J. Medvedovic, I. Bilic, M. Minnich, E. Axelsson, M. Jaritz, M. Busslinger, Essential role of EBF1 in the generation and function of distinct mature B cell types. Journal of Experimental Medicine 209, 775–792 (2012).

27. J.-H. Yang, J.-H. Li, P. Shao, H. Zhou, Y.-Q. Chen, L.-H. Qu, starBase: a database for exploring microRNA–mRNA interaction maps from Argonaute CLIP-Seq and Degradome-Seq data. Nucleic Acids Res 39, D202–D209 (2011).

28. J.-H. Li, S. Liu, H. Zhou, L.-H. Qu, J.-H. Yang, starBase v2.0: decoding miRNA-ceRNA, miRNA-ncRNA and protein–RNA interaction networks from large-scale CLIP-Seq data. Nucleic Acids Res 42, D92–D97 (2014).

29. D. W. Huang, B. T. Sherman, R. A. Lempicki, Bioinformatics enrichment tools: paths toward the comprehensive functional analysis of large gene lists. Nucleic Acids Res 37, 1–13 (2009).

30. D. W. Huang, B. T. Sherman, R. A. Lempicki, Systematic and integrative analysis of large gene lists using DAVID bioinformatics resources. Nature Protocols 4, 44– 57 (2009).

31. M. Kanehisa, S. Goto, KEGG: Kyoto Encyclopedia of Genes and Genomes. Nucleic Acids Res 28, 27–30 (2000).

32. M. Kanehisa, Y. Sato, M. Kawashima, M. Furumichi, M. Tanabe, KEGG as a reference resource for gene and protein annotation. Nucleic Acids Res 44, D457– D462 (2016).

33. M. Kanehisa, M. Furumichi, M. Tanabe, Y. Sato, K. Morishima, KEGG: new perspectives on genomes, pathways, diseases and drugs. Nucleic Acids Res 45, D353–D361 (2017).

34. H. S. Dengler, G. V. Baracho, S. A. Omori, S. Bruckner, K. C. Arden, D. H. Castrillon, R. A. DePinho, R. C. Rickert, Distinct functions for the transcription factor Foxo1 at various stages of B cell differentiation. Nature Immunology 9, 1388–1398 (2008).

35. T. Treiber, E. M. Mandel, S. Pott, I. Györy, S. Firner, E. T. Liu, R. Grosschedl, Early B Cell Factor 1 Regulates B Cell Gene Networks by Activation, Repression, and Transcription-Independent Poising of Chromatin. Immunity 32, 714–725 (2010).

36. H. Maier, R. Ostraat, H. Gao, S. Fields, S. A. Shinton, K. L. Medina, T. Ikawa, C. Murre, H. Singh, R. R. Hardy, J. Hagman, Early B cell factor cooperates with Runx1 and mediates epigenetic changes associated with mb-1 transcription. Nature Immunology 5, 1069–1077 (2004).

37. R. Song, P. Walentek, N. Sponer, A. Klimke, J. S. Lee, G. Dixon, R. Harland, Y. Wan, P. Lishko, M. Lize, M. Kessel, L. He, miR-34/449 miRNAs are required for motile ciliogenesis by repressing cp110. Nature 510, 115–120 (2014).

38. B. Vilagos, M. Hoffmann, A. Souabni, Q. Sun, B. Werner, J. Medvedovic, I. Bilic, M. Minnich, E. Axelsson, M. Jaritz, M. Busslinger, Essential role of EBF1 in the generation and function of distinct mature B cell types. Journal of Experimental Medicine 209, 775–792 (2012).

39. K. Hutter, T. Rülicke, M. Drach, L. Andersen, A. Villunger, S. Herzog, Differential roles of miR-15a/16-1 and miR-497/195 clusters in immune cell development and homeostasis. The FEBS Journal 288, 1533–1545 (2021).

40. M. Coffre, D. Benhamou, D. Rieß, L. Blumenberg, V. Snetkova, M. J. Hines, T. Chakraborty, S. Bajwa, K. Jensen, M. M. W. Chong, L. Getu, G. J. Silverman, R. Blelloch, D. R. Littman, D. Calado, D. Melamed, J. A. Skok, K. Rajewsky, S. B. Koralov, miRNAs Are Essential for the Regulation of the PI3K/AKT/FOXO Pathway and Receptor Editing during B Cell Maturation. Cell Reports 17, 2271– 2285 (2016).

41. H. Lin, R. Grosschedl, Failure of B-cell differentiation in mice lacking the transcription factor EBF. Nature 376, 263–267 (1995).

42. D. W. Harms, R. M. Quadros, D. Seruggia, M. Ohtsuka, G. Takahashi, L. Montoliu, C. B. Gurumurthy, Mouse Genome Editing Using the CRISPR/Cas System. Current Protocols in Human Genetics 83, 15.7.1–15.7.27 (2014).

43. C. A. Schneider, W. S. Rasband, K. W. Eliceiri, NIH Image to ImageJ: 25 years of image analysis. Nat Methods 9, 671–675 (2012).

44. M. R. Corces, A. E. Trevino, E. G. Hamilton, P. G. Greenside, N. A. Sinnott-Armstrong, S. Vesuna, A. T. Satpathy, A. J. Rubin, K. S. Montine, B. Wu, A. Kathiria, S. W. Cho, M. R. Mumbach, A. C. Carter, M. Kasowski, L. A. Orloff, V. I. Risca, A. Kundaje, P. A. Khavari, T. J. Montine, W. J. Greenleaf, H. Y. Chang, An improved ATAC-seq protocol reduces background and enables interrogation of frozen tissues. Nat Methods 14, 959–962 (2017).

