## Supplementary Figure 1-3 for "A single microRNA miR-195 rescues the arrested B cell development induced by EBF1 deficiency"

Fig. S1

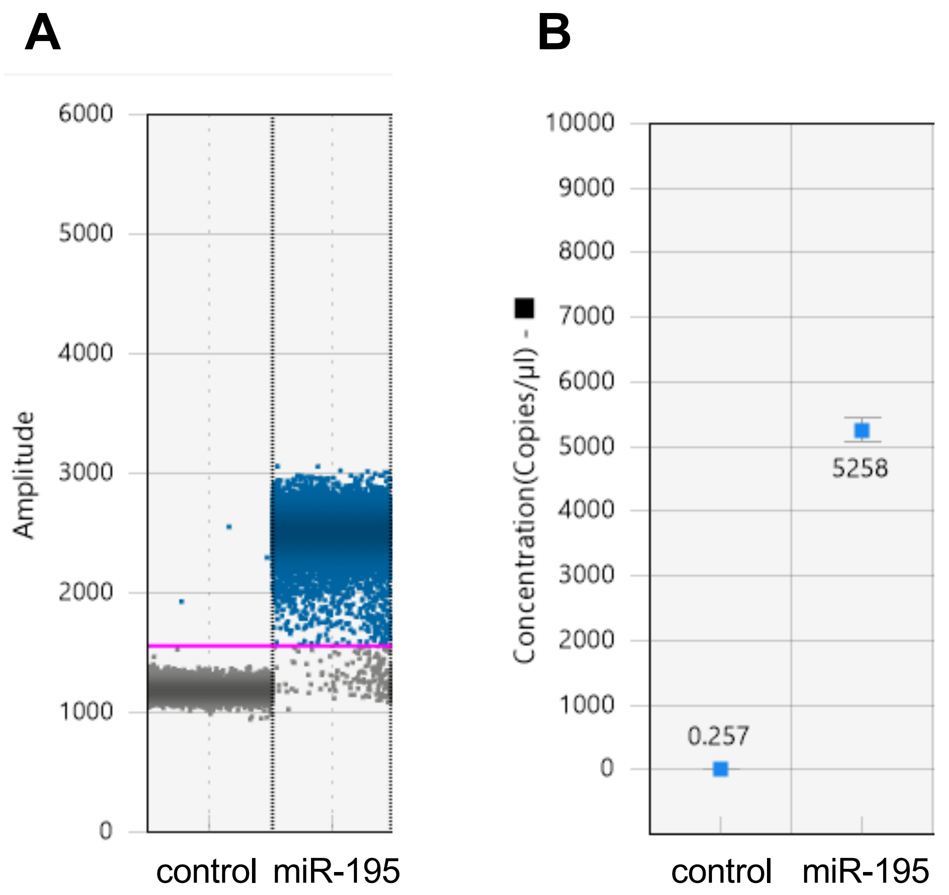


**Fig. S1. Raw data of Fig. 2C ddPCR.**
Amplitude (A) and concentration (B) of ddPCR on VJ recombination.

Fig. S2

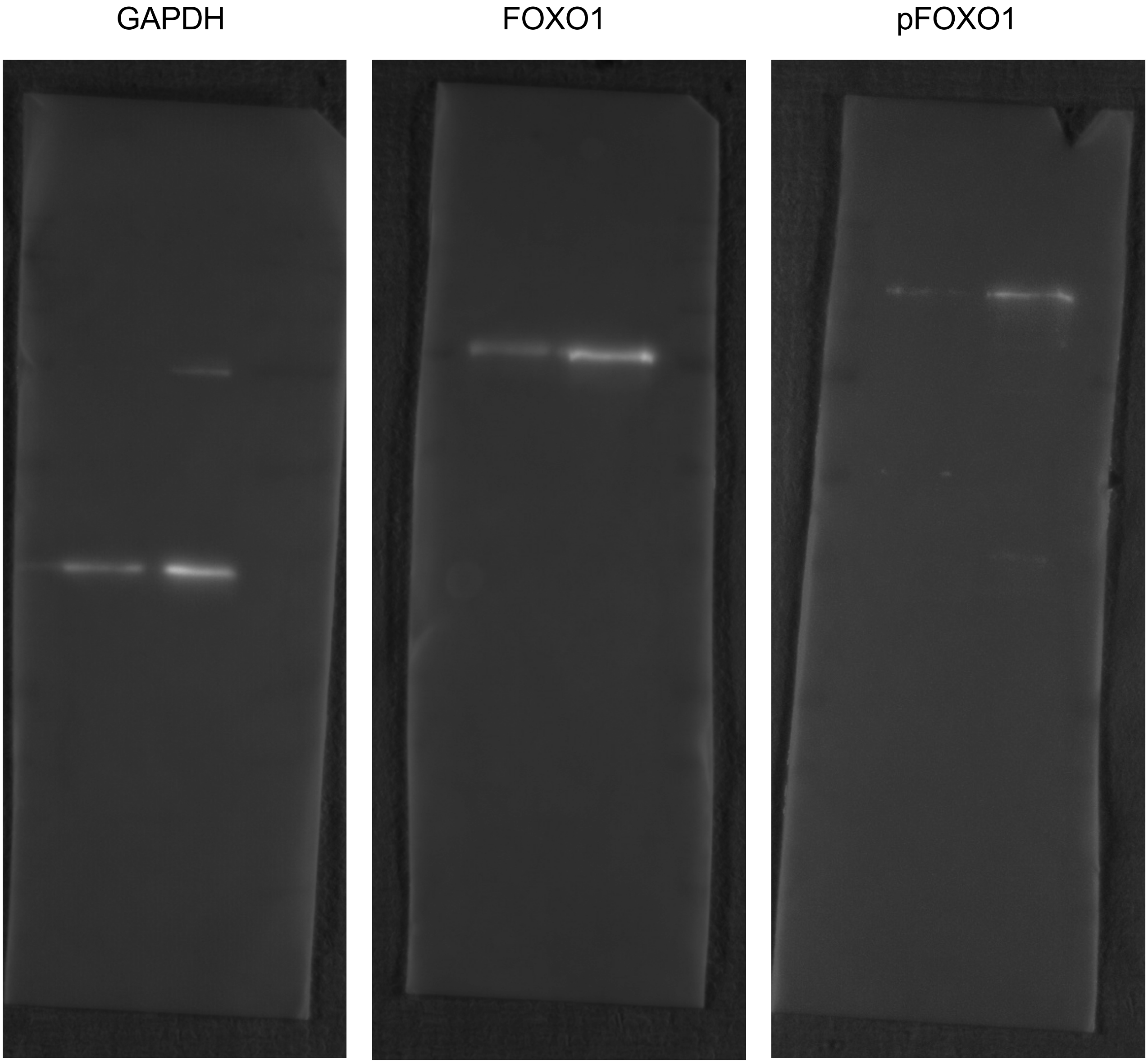


**Fig. S2. The uncropped Western blot images corresponding to Fig. 4C.**
miR-195 induced samples were placed in left side, and control samples were in right side. Precision Plus Protein™ standards (Bio-Rad Laboratories, Inc.) was used for markers.

### Fig. S3

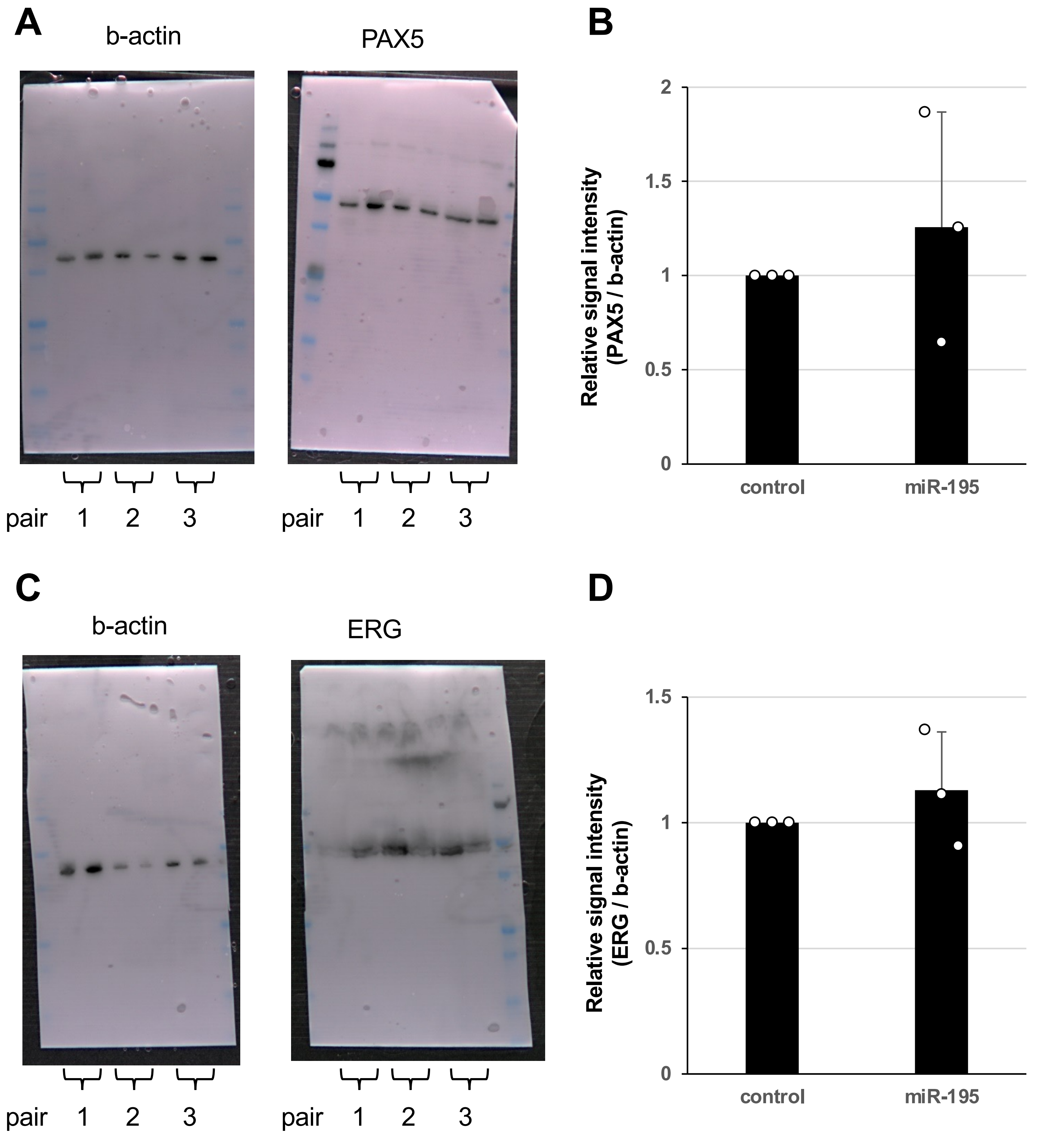


**Fig. S3. Western blot analysis of other known B-lineage regulators in Ebf1$^{-/-}$ HPCs with or without miR-195 transduction.**

(A-B) PAX5 and b-actin western blot images (A). 3 pairs of control (left) and miR-195 (right) transduced Ebf1$^{-/-}$ HPCs were analyzed. Quantification of b-actin and PAX5 band intensities from three independent samples is shown in the bar graph (B). Data are presented as mean ± SD. No significant difference in PAX5 protein level was observed.
(C-D) ERG and b-actin western blot images (C). ERG was analyzed same as PAX5, and Quantification of b-actin and ERG band intensities is also shown in the bar graph (D).
