## Supplementary Table 1-3 for "A single microRNA miR-195 rescues the arrested B cell development induced by EBF1 deficiency"

**Table S1. Lists of high-*p*-value pathways detected by means of targetome analysis of human miR-195 with target scan and starBase in the KEGG pathway.**

| Term | Count | P-Value | Benjamini |
|---|---|---|---|
| Neurotrophin signaling pathway | 38 | 2.20E-08 | 6.20E-06 |
| MAPK signaling pathway | 62 | 4.20E-08 | 5.80E-06 |
| Insulin signaling pathway | 40 | 1.30E-07 | 1.20E-05 |
| mTOR signaling pathway | 23 | 3.10E-07 | 2.20E-05 |
| Renal cell carcinoma | 24 | 7.20E-07 | 4.00E-05 |
| Acute myeloid leukemia | 21 | 3.20E-06 | 1.50E-04 |
| RNA transport | 42 | 8.00E-06 | 3.20E-04 |
| AMPK signaling pathway | 33 | 1.00E-05 | 3.50E-04 |
| Proteoglycans in cancer | 46 | 1.50E-05 | 4.50E-04 |
| HTLV-I infection | 55 | 1.50E-05 | 4.30E-04 |
| Pathways in cancer | 76 | 1.60E-05 | 4.10E-04 |
| PI3K-Akt signaling pathway | 68 | 2.60E-05 | 6.00E-04 |
| Signaling pathways regulating pluripotency of stem cells | 35 | 3.10E-05 | 6.70E-04 |
| Prostate cancer | 25 | 6.50E-05 | 1.30E-03 |
| Endocytosis | 53 | 8.00E-05 | 1.50E-03 |
| Insulin resistance | 28 | 1.20E-04 | 2.10E-03 |
| Focal adhesion | 44 | 1.50E-04 | 2.40E-03 |
| TNF signaling pathway | 27 | 2.20E-04 | 3.40E-03 |
| Regulation of actin cytoskeleton | 44 | 2.50E-04 | 3.70E-03 |
| Oocyte meiosis | 27 | 3.60E-04 | 4.90E-03 |
| Phosphatidylinositol signaling system | 25 | 3.90E-04 | 5.20E-03 |
| Progesterone-mediated oocyte maturation | 23 | 4.20E-04 | 5.30E-03 |
| Pancreatic cancer | 19 | 4.20E-04 | 5.10E-03 |
| Lysosome | 28 | 8.50E-04 | 9.80E-03 |
| FoxO signaling pathway | 30 | 9.30E-04 | 1.00E-02 |
| HIF-1 signaling pathway | 24 | 9.70E-04 | 1.00E-02 |
| Adherens junction | 19 | 1.30E-03 | 1.40E-02 |
| Ubiquitin mediated proteolysis | 30 | 1.30E-03 | 1.30E-02 |
| Wnt signaling pathway | 30 | 1.50E-03 | 1.40E-02 |
| Hippo signaling pathway | 32 | 1.60E-03 | 1.50E-02 |
| Epstein-Barr virus infection | 38 | 1.60E-03 | 1.40E-02 |
| Non-small cell lung cancer | 16 | 1.80E-03 | 1.60E-02 |
| mRNA surveillance pathway | 22 | 2.00E-03 | 1.70E-02 |
| Ras signaling pathway | 43 | 2.10E-03 | 1.70E-02 |
| Cell cycle | 27 | 2.70E-03 | 2.10E-02 |
| Shigellosis | 17 | 2.80E-03 | 2.20E-02 |
| Hepatitis B | 30 | 3.30E-03 | 2.50E-02 |
| Choline metabolism in cancer | 23 | 3.40E-03 | 2.50E-02 |
| Prolactin signaling pathway | 18 | 3.40E-03 | 2.40E-02 |
| Chronic myeloid leukemia | 18 | 4.00E-03 | 2.80E-02 |
| Small cell lung cancer | 20 | 4.60E-03 | 3.10E-02 |

**Table S2. Lists of high-*p*-value pathways detected by means of targetome analysis of murine miR-195 with target scan and starBase in the KEGG pathway.**

| Term | Count | P-Value | Benjamini |
|---|---|---|---|
| Insulin signaling pathway | 18 | 5.60E-06 | 1.40E-03 |
| Protein processing in endoplasmic reticulum | 19 | 1.70E-05 | 2.10E-03 |
| Focal adhesion | 21 | 2.80E-05 | 2.30E-03 |
| Pancreatic cancer | 11 | 6.70E-05 | 4.10E-03 |
| PI3K-Akt signaling pathway | 28 | 6.90E-05 | 3.40E-03 |
| Neurotrophin signaling pathway | 15 | 7.20E-05 | 2.90E-03 |
| Cell cycle | 15 | 8.60E-05 | 3.00E-03 |
| Toxoplasmosis | 14 | 1.30E-04 | 3.90E-03 |
| TGF-beta signaling pathway | 12 | 1.50E-04 | 4.00E-03 |
| Prostate cancer | 12 | 2.00E-04 | 4.90E-03 |
| Signaling pathways regulating pluripotency of stem cells | 15 | 2.70E-04 | 6.00E-03 |
| Colorectal cancer | 10 | 3.10E-04 | 6.30E-03 |
| Renal cell carcinoma | 10 | 4.40E-04 | 8.30E-03 |
| Phosphatidylinositol signaling system | 12 | 4.70E-04 | 8.20E-03 |
| Pathways in cancer | 28 | 5.30E-04 | 8.50E-03 |
| HTLV-I infection | 22 | 5.80E-04 | 8.80E-03 |
| Small cell lung cancer | 11 | 5.80E-04 | 8.30E-03 |
| FoxO signaling pathway | 14 | 6.90E-04 | 9.30E-03 |
| Progesterone-mediated oocyte maturation | 11 | 7.70E-04 | 9.80E-03 |
| Prolactin signaling pathway | 10 | 8.40E-04 | 1.00E-02 |
| Hepatitis B | 14 | 1.50E-03 | 1.80E-02 |
| Endometrial cancer | 8 | 1.90E-03 | 2.10E-02 |
| Hippo signaling pathway | 14 | 2.10E-03 | 2.20E-02 |
| Measles | 13 | 2.50E-03 | 2.50E-02 |
| Hepatitis C | 13 | 2.50E-03 | 2.50E-02 |
| Non-small cell lung cancer | 8 | 2.90E-03 | 2.80E-02 |
| HIF-1 signaling pathway | 11 | 3.00E-03 | 2.80E-02 |
| Chronic myeloid leukemia | 9 | 3.20E-03 | 2.80E-02 |
| Regulation of lipolysis in adipocytes | 8 | 3.20E-03 | 2.80E-02 |
| Ubiquitin mediated proteolysis | 13 | 3.50E-03 | 2.90E-02 |
| Sphingolipid signaling pathway | 12 | 3.60E-03 | 2.90E-02 |
| AMPK signaling pathway | 12 | 4.30E-03 | 3.30E-02 |
| Epstein-Barr virus infection | 16 | 7.30E-03 | 5.40E-02 |
| Acute myeloid leukemia | 7 | 1.20E-02 | 8.60E-02 |
| Oocyte meiosis | 10 | 1.30E-02 | 9.20E-02 |
| Endocytosis | 18 | 1.50E-02 | 1.00E-01 |
| Rap1 signaling pathway | 15 | 1.60E-02 | 1.00E-01 |

**Table S3. List of genes with detected peaks in the DNA region common in the three analyses.**

| Gene names having open chromatin peak(s) in WT $CD19^{+}$ vs EBF1 KO $CD19^{-}$ and WT $CD19^{+}$($\sim IgM^{+}$) vs WT $CD19^{-}$ and miR-195 $CD19^{+}$ vs EMPTY $CD19^{-}$ |
|---|
| Ablim1, Acoxl, Alk, Ankrd6, Atg7, B230118H07Rik, BC003331, Bfsp1, Blnk, Cacna1d, Capn11, Cdc42se2, Ciita, Cmip, Dennd4a, Dio2, Ebf1, Eif2ak3, Entpd5, Epb41l2, Erg, Foxn3, Foxp1, Gabrr2, Gpr20, Grik4, Hivep3, Irf8, Itga9, Itpa, Itpr2, Jakmip1, Kalrn, Ldlrad4, Lipc, Med13l, Mgll, Mical3, Myo1d, Nav2, Nfia, Nsmce2, Osbpl10, Pakap, Pax5, Pde2a, Pde4c, Pgm5, Plxnc1, Pold1, Prlr, Rarres1, Rassf5, Rd3, Rnf41, Runx1, Ryr2, Scn4a, Sh3bp5, Slc39a11, Spink2, Stxbp1, Tbc1d1, Tg, Thada, Tmem64, Trappc9, Trio, Ubac2, Ube2h, Vps8, Whrn, 2410004B18Rik |
